# Hippocampal OLM interneurons regulate CA1 place cell plasticity and remapping

**DOI:** 10.1101/2024.11.11.622941

**Authors:** Matt Udakis, Matthew Claydon, Heng Wei Zhu, Jack R Mellor

## Abstract

OLM interneurons selectively target inhibition to the distal dendrites of CA1 pyramidal cells in the hippocampus but the role of this unique morphology in controlling place cell physiology remains a mystery. Here we show that OLM activity prevents associative synaptic plasticity at Schaffer collateral synapses on CA1 pyramidal cells by inhibiting dendritic Ca^2+^ signalling initiated by entorhinal synaptic inputs. Furthermore, we find that OLM activity is reduced in novel environments suggesting that reducing OLM activity and thereby enhancing excitatory synaptic plasticity is important for the formation of new place cell representations. Supporting this, we show that selectively increasing OLM activity in novel environments enhances place cell stability and reduces remapping of newly formed place cells whilst increasing OLM activity in familiar environments led only to a transient silencing of place cells. Our results therefore demonstrate a critical role for distal dendrite targeting interneurons in regulating plasticity of neuronal representations.

## Introduction

The hippocampus enables us to learn and adapt to new situations by maintaining a dynamic representation of the environment. Location specific place cell activity allows quantifiable measurement of the neuronal representation of an environment and when a new environment is encountered, hippocampal place cells rapidly adapt their firing (or remap) to efficiently encode the new features of the environment ^1–4^. This dynamic neuronal representation is essential for behavioural adaptation to new environments ^5^. Synaptic plasticity triggered by Ca^2+^ influx to dendrites and spines is a fundamental process that underpins adaptation of place cell firing where large dendritic Ca^2+^ events termed plateau potentials have been shown to be particularly effective at driving the formation and tuning of place cells ^6–13^. In CA1 pyramidal cells, plateau potentials are thought to be driven by supra-linear integration of coincident CA3 (proximal) and entorhinal (distal) inputs ^6, 7, 9, 14, 15^, but the regulation of this integration, and therefore of behaviorally relevant synaptic plasticity and place cell adaptation, is largely unexplored.

Hippocampal interneurons in CA1 are diverse and sub-categorised by genetic, morphological and physiological properties ^16–20^. Major sub-categories are parvalbumin (PV) expressing interneurons that are mostly perisomatic targeting and somatostatin (SOM) expressing interneurons that are mostly dendrite targeting ^17, 19^, but within these general subtypes there is considerable heterogeneity ^21^. Manipulation of all CA1 interneuron activity disrupts many features of place cell activity including place cell formation ^22–24^, specifically, targeted silencing of PV interneurons alters the timing of place cell firing with respect to theta cycles whereas SOM interneuron silencing enhances place cell burst firing ^25^. The dendritic targeting of inhibition by SOM interneurons imparts them with privileged control of synaptic inputs whereas perisomatic targeting of inhibition by PV interneurons provides them with fine control of pyramidal cell spike timing and rate ^26^. This suggests that SOM interneurons are prime candidates to regulate synaptic integration, plateau potentials, and thus the plasticity associated with place cell formation and remapping.

Activity of different interneuron subtypes is also associated with behavioural state induced by experience. In novel environments, where plasticity enables place cells to remap to the new environment, interneurons increase or decrease their activity in line with their subtype: PV interneurons generally increase activity and SOM interneurons decrease firing rate, but with considerable variability perhaps reflecting distinct subtypes within the PV or SOM classification (see ^27–31^).

Within the family of CA1 SOM interneurons, Oriens Lacunosum Moleculare (OLM) interneurons have a unique morphology receiving feedback excitation from CA1 in stratum oriens and targeting inhibition specifically to the distal dendrites of CA1 pyramidal cells in stratum lacunosum moleculare. They also receive direct cholinergic input from the medial septum and express nicotinic and muscarinic receptors that respond to the release of acetylcholine ^32–35^. Indeed, the majority of OLM neurons express the α2 nicotinic receptor (OLMα2; ^35, 36^), which can be used to selectively manipulate and measure OLMα2 neurons using Chrna2-cre mice ^33^. Surprisingly, stimulation of OLMα2 neurons in *ex vivo* slices enhances synaptic plasticity at CA3-CA1 Schaffer collateral synapses via a disinhibitory action on PV interneurons ^33^ but the role of OLMα2 neurons in regulating plasticity driven by entorhinal inputs and plateau potentials has not been investigated. Furthermore, disruption of OLMα2 activity impairs learning in novel object recognition, y-maze and fear conditioning paradigms ^34, 37–39^ but the contribution of OLM neurons to the plasticity of neuronal representations required for hippocampal-dependent learning is unknown.

Here we made use of Chrna2-cre mice to selectively measure and manipulate OLMα2 interneurons in the hippocampus. By optogenetically stimulating OLM interneurons in *ex vivo* slices we show that they control dendritic Ca^2+^ signalling and synaptic plasticity initiated by entorhinal inputs to CA1 pyramidal neurons. Measurement of OLM activity during behaviour revealed a strong regulation by novelty and movement where OLM firing was reduced in a novel environment that requires place cell remapping. Finally, bidirectional manipulation of OLM activity at specific locations within novel and familiar environments showed that OLM interneurons regulate the emergence and stability of new place cell representations. Together, these results demonstrate a critical role for dendrite targeted inhibition for control of synaptic plasticity that underpins the remapping of neuronal representations necessary for adapting to unfamiliar environments.

## Results

### Control of synaptic plasticity and dendritic calcium dynamics by OLM interneurons

Behaviorally relevant synaptic plasticity that underlies place cell adaptations in CA1 is thought to be induced by coincident CA3 and entorhinal cortex inputs that summate supra-linearly and generate large dendritic Ca^2+^ events termed plateau potentials ^6, 7, 9, 14, 15^. The precise targeting of OLM interneuron synapses to the distal dendrites of CA1 pyramidal cells where entorhinal cortex inputs are located suggests they may have an important role in the integration of entorhinal and CA3 inputs and therefore generation of behaviorally relevant synaptic plasticity. To test this we selectively activated a subset of OLM interneurons that express Chrna2 ^33, 36^ by expressing the light-activated cation channel channelrhodopsin-2 (ChR2) in a cre-dependent manner using mice that expressed cre recombinase under control of the promoter for Chrna2 (Chrna2-cre) crossed with mice expressing cre-dependent ChR2 (Ai32 mice; methods). Immunohistochemistry confirmed that Chrna2-cre mice expressed cre selectively in OLM interneurons with reporter expression highest in the Stratum Oriens (SO) and Stratum Lacunosum Moleculare (SLM) layers and cell bodies principally located in SO (Figure 1A) ^33,40^. This expression profile is consistent with OLM interneurons providing distal dendritic inhibition that exhibits slower synaptic current kinetics when measured at the soma than perisomatic targeting parvalbumin expressing interneurons ^41^.

**Figure 1.**
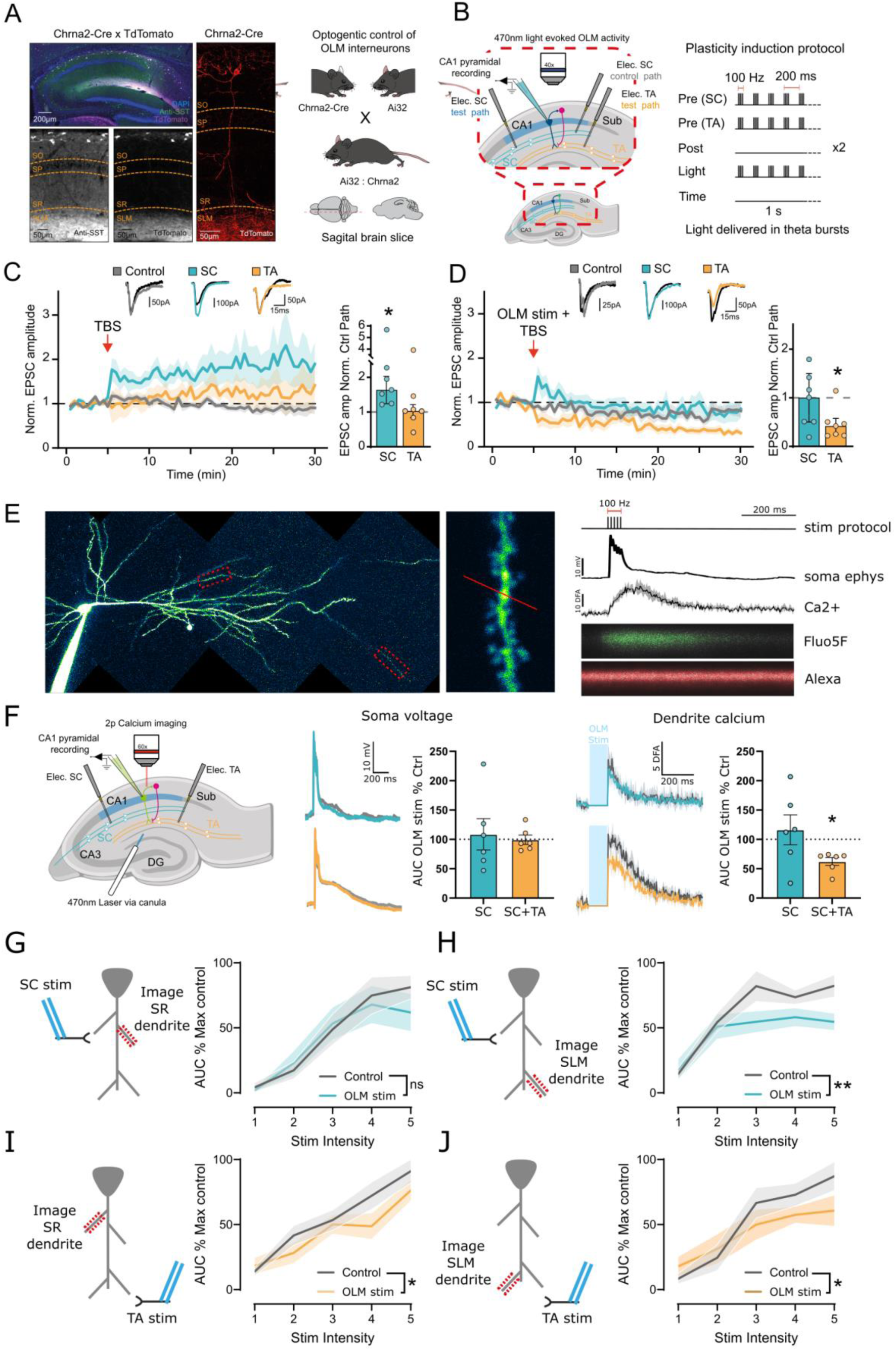
– OLM neurons regulate associative synaptic plasticity and dendritic Ca^2+^. (A) Immunohistochemistry showing expression of SST and tdTomato in Chrna2-cre x loxP-stop-loxP-tdTomato mice in different hippocampal layers: Stratum Oriens (SO), Stratum Pyramidal (SP) Stratum Radiatum (SR) and Stratum Lacunosum Moleculare (SLM). (B) Schematic of experimental approach. *Ex vivo* hippocampal slices prepared from Chrna2-cre x loxP-stop-loxP-ChR2 mice and recordings made from CA1 pyramidal cells whilst stimulating Schaffer collateral (SC), temporoammonic (TA) and OLM inputs. Plasticity protocol shown on right with or without light stimulation of OLM interneurons. (C) Theta Burst Stimulation (TBS) delivered concurrently to SC and TA inputs induced LTP at test pathway SC but not TA synapses. Example traces before and after plasticity shown above (P < 0.05, paired t-test, two-tailed, n = 7 cells). (D) Same as C but with light stimulation of OLM interneurons during TBS. No LTP induced in SC test pathway and LTD induced in TA pathway (P < 0.05, paired t-test, two-tailed, n = 7 cells). (E) CA1 pyramidal cell filled with Alexa594 and Fluo5. Recorded SR and SLM regions of dendrite are highlighted with red dashed boxes with high magnification image of line scan region (middle). Left, example recording from illustrated line scan with dF/F quantified trace and somatically recorded voltage in response to concurrent stimulation of SC and TA inputs. (F) OLM stimulation reduces SR dendritic Ca^2+^ in response to concurrent TA and SC inputs but not when only SC inputs are stimulated. Example somatic EPSPs and SR dendrite Ca^2+^ responses are shown with and without OLM stimulation. Note blanked region of Ca^2+^ response during light stimulation of OLM interneurons (P < 0.05, paired t-test, two-tailed, n = 6). (G,H,I,J) Increasing stimulation intensity to SC (G,H) or TA (I,J) input pathways increases Ca^2+^ responses in SR (G,I) or SLM (H,J) dendritic regions. OLM stimulation reduces dendritic Ca^2+^ responses in H,I&J but not G (P < 0.05 in H,I,J, Two-way ANOVA, n = 6,7,6,6 For G,H,I,J respectively). Data presented as mean values ± S.E.M.

Having established selective control of OLM interneurons we investigated whether OLM interneurons can control behaviorally relevant synaptic plasticity. Excitatory postsynaptic currents (EPSCs) elicited in 3 independent pathways (Schaffer collateral (SC) control, SC test and temporoammonic (TA) test) were recorded from CA1 pyramidal neurons, via whole-cell patch clamp. Plateau potentials can be replicated in *ex vivo* slices by bursts of high frequency synaptic input ^7, 42, 43^, therefore, to induce synaptic plasticity, coincident theta burst stimulation (TBS) was given to the SC and TA test pathways. Since OLM interneurons are feedback interneurons excited by CA1 pyramidal neurons and active during the same phase of theta cycles *in vivo* ^17, 18^, the impact of OLM input was assessed by concurrent TBS of OLM interneurons (Figure 1B). In the absence of OLM stimulation LTP was induced in the SC test pathway but not in the TA test pathway (Figure 1C; 2.22 ± 0.59 (SC), 1.39 ± 0.44 (TA), n = 7). In contrast, in the presence of OLM stimulation SC LTP was blocked and instead LTD emerged in the TA test pathway (Figure 1D; 0.94 ± 0.22 (SC), 0.46 ± 0.12 (TA), n = 7). This indicates that when synaptic plasticity is induced by a behaviorally relevant synaptic stimulation, OLM interneurons inhibit LTP at SC synapses.

Plateau potentials in CA1 pyramidal neurons generate large Ca^2+^ increases in SR apical dendrites which triggers synaptic plasticity at SC synapses, but OLM interneurons target SLM distal dendrites. Therefore, we next tested whether OLM interneurons can regulate Ca^2+^ in SR dendrites during plateau potentials. To measure dendritic Ca^2+^ fluctuations, Ca^2+^ dynamics were imaged in *ex vivo* slices using 2photon imaging of defined apical SR and SLM dendritic regions during electrical synaptic stimulation of SC and/or TA inputs. CA1 pyramidal neurons were filled with the morphological marker Alexa594 and the Ca^2+^ indicator Fluo5f and dendritic Ca^2+^ and somatic voltage responses to synaptic stimulation recorded (Figure 1E). Coincident TBS of TA and SC inputs that induced LTP in SC synapses generated large depolarizations at the soma that lasted for an extended period (>200ms) similar to plateau potentials that induce new place fields *in vivo* ^6–13^. These depolarizations were associated with large and long-lasting dendritic Ca^2+^ increases in the SR region of apical dendrites (Figure 1F). Concurrent TBS of OLM interneurons substantially reduced the SR dendritic Ca^2+^ signal without any impact on the somatically recorded depolarization (Figure 1F; 62.28 ± 6.81 % (Ca^2+^), 99.41 ± 8.01 % (depolarization)). This is reminiscent of other situations where dendritic and synaptic voltage dynamics are localised and do not propagate linearly to the soma^15, 44, 45^. In comparison, when SC inputs were stimulated in the absence of TA stimulation, smaller depolarizations and Ca^2+^ increases were generated and no effect of coincident OLM interneuron stimulation was observed (Figure 1F; 116.3 ± 25.59 % (Ca^2+^), 108.6 ± 26.69 % (depolarization)). This indicates that OLM interneurons can reduce the Ca^2+^ response in SR dendrites by preventing the integration of coincident TA inputs.

To further test this conclusion, we spatially dissected the action of OLM interneurons on dendritic Ca^2+^ by stimulating SC and TA inputs separately and measuring the impact of OLM interneuron stimulation on dendritic Ca^2+^ in apical SR and SLM regions. In these experiments dendritic Ca^2+^ increases were induced by increasing the stimulation intensity for each input pathway in a step wise manner. OLM stimulation decreased Ca^2+^ transients in the distal SLM dendrites regardless of the site of synaptic stimulation whereas Ca^2+^ transients were only decreased at proximal SR dendrites when TA inputs were stimulated and not when SC inputs were stimulated alone (Figure 1G,H,I,J; SLM/SC – F(1, 6) = 20.46, P = 0.004, SLM/TA – F(1, 5) = 8.298, P = 0.0346, SR/TA – F(1, 5), P = 0.0201, SR/SC – F(1, 5) = 0.389, P = 0.5602). This demonstrates that OLM interneurons control not only the Ca^2+^ transients at distal dendrites but also Ca^2+^ transients in proximal dendrites if the depolarization originates in distal dendrites. This provides an explanation for how OLM interneurons control SR dendrite Ca^2+^ during plateau potentials and therefore synaptic plasticity at SC synapses on apical CA1 dendrites.

### OLM activity during exploration in novel environments

Synaptic plasticity leading to place cell formation and remapping is predicted to occur when mice explore novel environments. The question we next addressed is whether OLM interneurons play a role in enabling synaptic plasticity in novel environments by altering their firing rate. To answer this question, OLM activity was monitored *in vivo* by measuring Ca^2+^ dynamics in OLM interneurons using miniaturised microscopy during exploration of familiar and novel environments. AAV1-Flex-GCaMP6s was injected into the dorsal hippocampus of Chrna2-cre mice and a GRIN lens and baseplate implanted above the hippocampus to selectively express and image the Ca^2+^ indicator GCaMP6s in OLMα2 interneurons (Figure 2A). Mice were trained to run from end to end of a 140cm linear track for 10% sucrose rewards whilst tethered to a miniscope camera that recorded the activity of OLM interneurons (Figure 2B). To investigate the response of these interneurons to novel environments OLM activity was first observed during exploration of a familiar environment for 10 minutes before switching the mice to a novel environment. The average OLM activity on the initial laps exploring a novel environment was significantly lower than the activity of the latter laps of exploration in a familiar environment (0.011 ± 0.005 vs 0.093 ± 0.063 DF/F). In comparison, mice transitioned between two exposures of the familiar environment showed no change in OLM activity (0.084 ± 0.008 vs 0.082 ± 0.008 DF/F) (Figure 2 C,D). Analysis of single interneurons revealed the majority of interneurons reduced their activity in the novel environment and this reduction was more prominent for interneurons that were more active in the familiar environment (Figure 2E). As expected during novel exploration the velocity of the mice was reduced, as the mice explored unfamiliar surroundings (Figure 2F) and similar to other interneuron populations within the hippocampus ^29, 31, 46^ we observed that OLM activity was also correlated with velocity of the mice (Figure 2B,G,H). Therefore, it was important to distinguish between any reduction in OLM activity due to a change in velocity and changes in OLM activity due to a novelty signal. To do this the average activity of the interneurons was binned based on the velocity of the mice. This revealed that at comparable velocities the activity of OLM interneurons was consistently reduced (Figure 2G). The correlation between interneuron activity and mouse velocity was still observed in the novel environment, although significantly reduced compared to familiar exploration (Figure 2G,H). These observations suggest that OLM activity is positively modulated by mouse velocity and negatively modulated by novelty suggesting that synaptic plasticity at CA1 pyramidal neurons is more likely to occur in novel environments when place cells are remapped.

**Figure 2.**
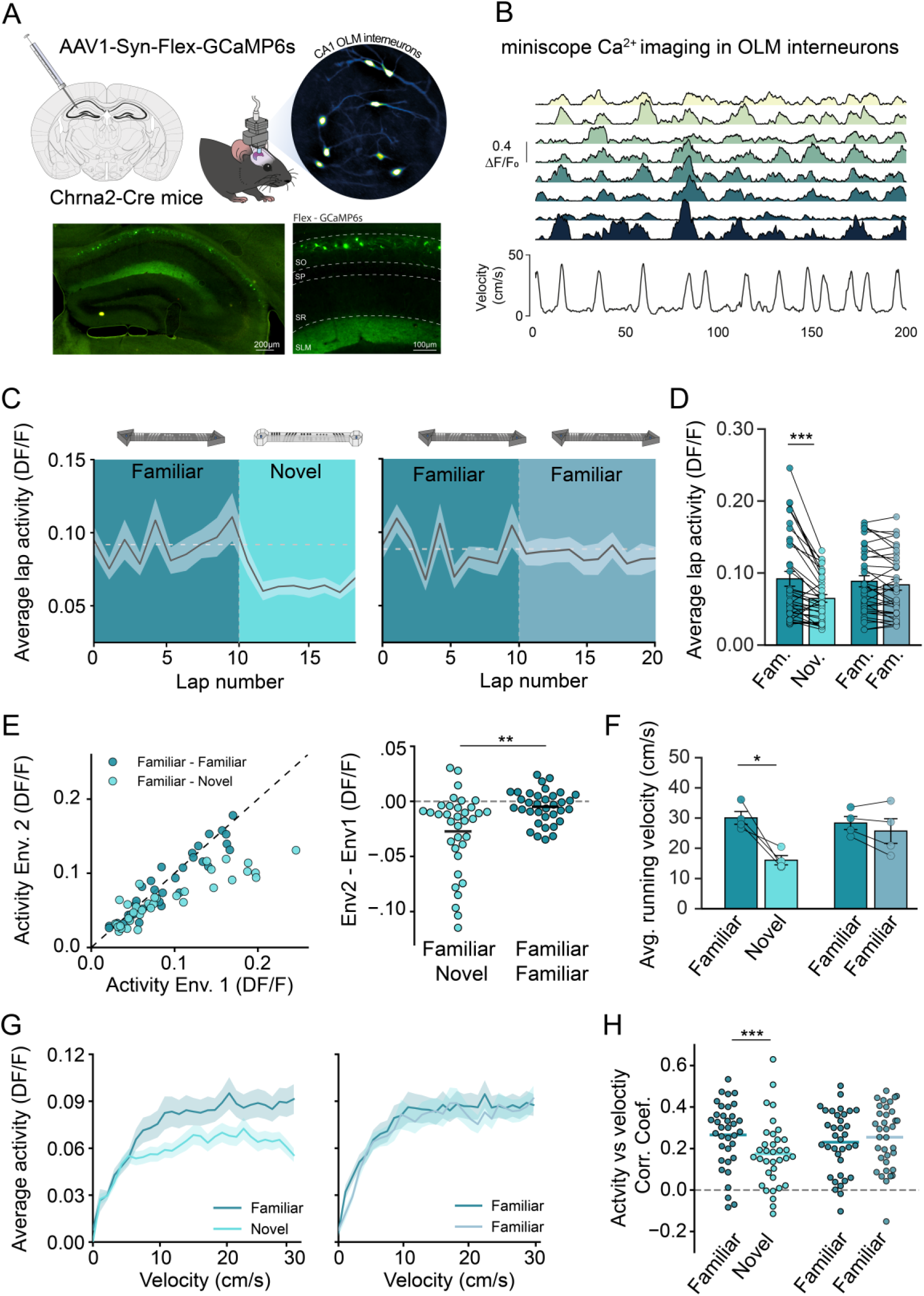
– OLM activity reduces in novel environments. (A) Chrna2-cre mice were injected in the dorsal hippocampus with a CRE dependent viral construct to express GCaMP6s in OLM interneurons. Miniscope field of view of GCaMP6s expression in OLM interneurons. Widefield fluorescence of dorsal hippocampus brain slice showing expression of GCaMP6s restricted to OLM interneurons. (B) Example Ca^2+^ fluorescence recording from 8 OLM interneurons within the same mouse together with the corresponding mouse velocity as the mouse explored a familiar environment. (C) Average lap-by-lap activity of OLM interneurons as mice explored a familiar and then novel environment (left) and two consecutive familiar environments (right). (D) Average activity of individual OLM interneurons as they explored different environments. Average activity across all laps n = 35 cells from 4 mice. Paired t-test ***: *p* < 0.001. (E) Average activity of each interneuron in familiar environment vs average activity in the subsequent environment (novel or familiar) (left). Activity difference between the two environments. Paired t-test **: *p* < 0.01. (F) Average running velocity of mice during exploration of familiar and novel environments. n = 4 mice, paired t-test *: *p* < 0.05. (G) Average OLM interneuron activity at different mouse velocities during exploration of familiar and novel environments. (H) OLM activity vs mouse velocity Pearsons correlation coefficients between familiar and novel environments and two consecutive exposures to a familiar environment. Paired t-test ***: *p* < 0.001. Data presented as mean values ± S.E.M.

### Place cell formation and remapping is modulated by OLM interneuron activity

With OLM interneurons reducing their activity in novel environments and able to modulate excitatory synaptic plasticity we hypothesised that OLM interneurons may play a key role in the mechanisms of place cell formation and the plasticity that leads to their long-term stability^47^. To test this, we recorded place cell formation and stability *in vivo* whilst manipulating the activity of OLM interneurons using miniaturised microscopy. To express the Ca^2+^ indicator GCaMP6f into dorsal hippocampal CA1 pyramidal neurons, we injected Chrna2-cre mice with AAV5-CAMKII-GCaMP6f alongside either AAV5-hSyn-Flex-ChrimsonR, AAV5-hSyn-Flex-JAWs or the control AAV5-hsyn-DIO-mCherry to express either the redshifted excitatory opsin ChrimsonR or the redshifted inhibitory opsin JAWs in OLMα2 interneurons. This enabled us to record the activity of pyramidal neurons whilst either stimulating or inhibiting the activity of OLM interneurons. OLM activity was modulated using 620 nm light through the miniscope when mice entered a specific 33 cm section of the track termed the ‘optogenetic-zone’ with an adjacent 33 cm section of the track without light delivery used as a direct comparator termed the ‘control zone’ (Figure 3A).

**Figure 3.**
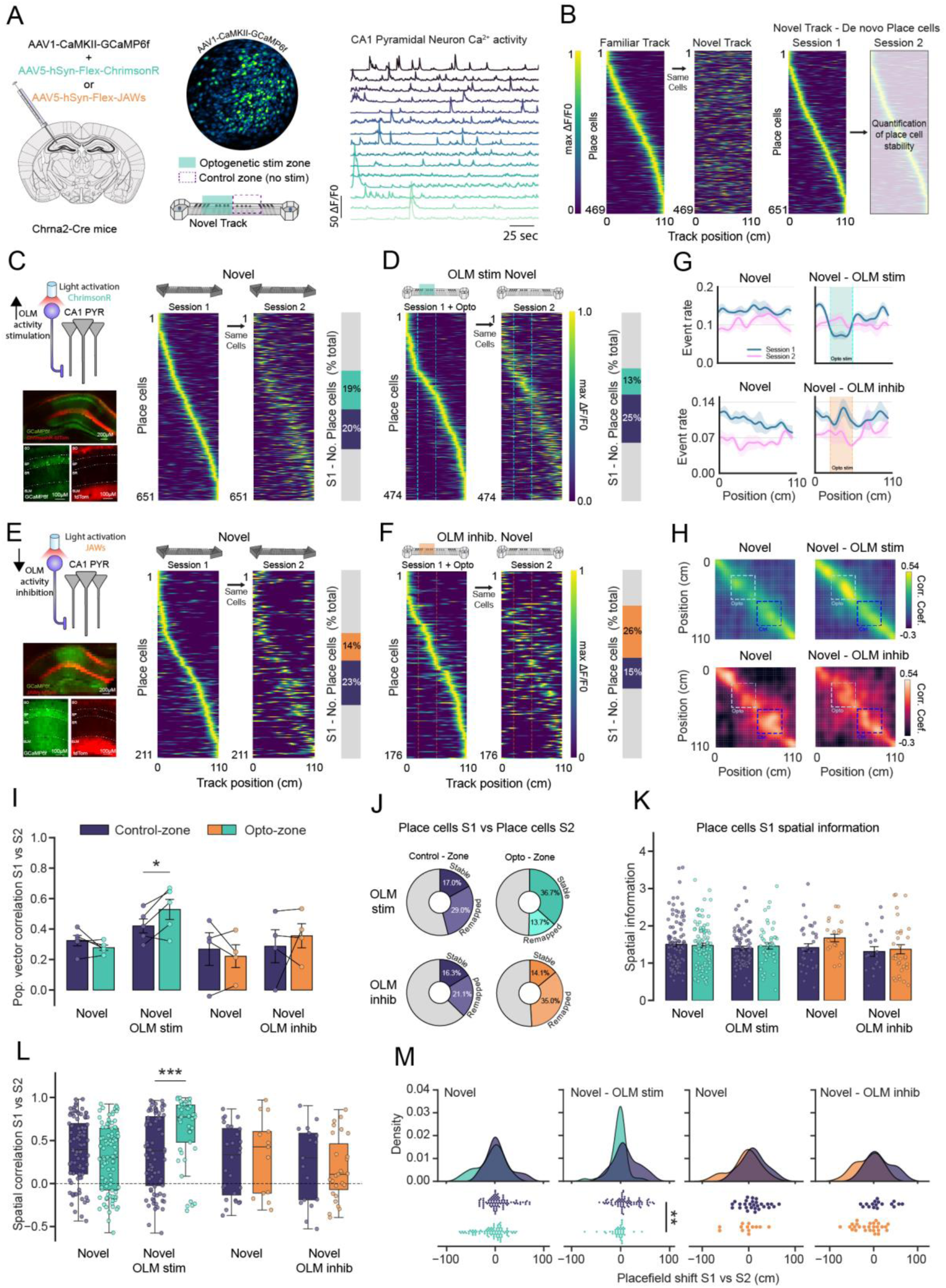
– OLM activity regulates stability of spatial representations in novel environments. (A) Chrna2-cre mice were injected into the dorsal hippocampus with viral constructs to express GCaMP6f in pyramidal and either ChrimsonR or JAWs into OLM interneurons (left). Miniscope field of view of GCaMP6f expression in pyramidal neurons and example Ca^2+^ transients from recorded pyramidal neurons. Opsins were activated at a specific portion of the track (optogenetic stim zone), data were then compared to a control zone. (B) Place cell firing rate maps in a familiar environment with the same cells firing rates in a novel environment, demonstrating global remapping in a novel environment. *De novo* place cells were acquired during the exploration of the novel environment that tiled the full length of the track. (C) Example expression of excitatory opsin ChrimsonR in OLM interneurons to increase the activity of OLM interneurons (left). *De novo* place cells acquired during the first exposure to the novel environment (session 1) in the absence of optogenetic stimulation. Activity of the same cells in session 1 during the second exposure to the novel environment (session 2) (middle). Proportions of place cells during session 1 encoding different zones on the track. (D) *De novo* place cells acquired in first exposure to a novel environment (session 1) with optogenetic activation of ChrimsonR to stimulate OLM interneuron activity. Activity of the same cells in session 1 during the second exposure to the novel environment (session 2) (left). Proportions of place cells during session 1 encoding different zones on the track. (E) Same as C except for expression of JAWs opsin to decrease activity in OLM interneurons. (F) Same as D except for optogenetic activation of JAWs to decrease OLM interneuron activity. (G) Ca^2+^ event rate of all recorded neurons along the length of the track during the exploration of novel environments (OLM stim (top row) n = 5 mice, OLM inhib. (bottom row) n = 4 mice). (H) Population vector correlation matrix between activity rate maps in session 1 vs session 2 for C-F, averaged across animals. (I) Average population vector correlation taken as average diagonal correlation for each track zone, (Control-zone vs Opto-zone). (OLM stim n = 5 mice, OLM inhib. n = 4 mice, (*: *p* < 0.05, paired t-test). (J) Retention of session 1 place cells during session 2, place cells were either lost, and no longer classified as place cells (grey), stable places encoding a similar location in session 2 or remapped place cells encoding a new location in session 2. Average % across animals. (K) Spatial information content of session 1 place cells in the control and optogenetic zones. (L) Spatial correlation values between place cells in session 1 and the activity of the same cells in session 2, for both control and optogenetic zones. Unpaired t-test on Fisher corrected data ***: *p* < 0.001 (M) Place field location shifts from place cells in session 1 vs place field location of same cells in session 2. Unpaired t-test on absolute position shifts **: *p* < 0.01. Data presented as mean values ± S.E.M.

Analysis of CA1 pyramidal neuron Ca^2+^ activity whilst mice traversed a familiar linear track revealed place cell firing fields that tiled the entire length of the track. When mice were switched to a novel track the spatial encoding of these cells was completely lost and a new spatial representation was formed from a population of *de novo* place cells. (Figure 3B). To assess the impact of OLM activity on place cell formation and remapping, pyramidal neuron activity was recorded during exploration in a novel environment (session 1) with and without optogenetic manipulation of OLM interneurons. To subsequently assess the stability of these newly formed place cells and the impact of OLM perturbation, mice were re-exposed to the same novel environment for a second time (session 2) and place cell representations were compared between Session 1 and Session 2.

Increased OLM activity by activation of chrimsonR in the optogenetic stimulation zone of the novel track reduced the proportion of *de novo* place cells that represented that spatial location (Figure 3C,D). This corresponded with a reduction in the event rate of the pyramidal neurons specific to the region of the track where OLM activity was increased. Together with the lack of any effects of LED stimulation in the mCherry expressing mice (Figure S1), this confirms the predicted impact of enhanced inhibition on pyramidal neuron firing and demonstrates the successful implementation of OLM interneuron optogenetic excitation (Figure 3G). In contrast, reducing the activity of OLM interneurons with the inhibitory opsin JAWs caused a small and non-significant increase of pyramidal neuron activity that lead to an overrepresentation of *de novo* place cells selective to the optogenetic stimulation zone (Figure 3E,F).

To assess the stability of newly formed spatial representation we calculated the population vector correlations between the two exposures to the same novel environment (session 1 vs session 2) This revealed that increasing OLM activity led to enhanced spatial encoding restricted to the optogenetic stimulation zone (0.421 ± 0.046 vs 0.529 ± 0.066 (correlations for control vs opto zone)). Interestingly, reducing the activity of OLM interneurons did not lead to a decrease in the population encoding of the track (Figure 3H,I).One possible explanation for enhanced spatial encoding between the two sessions would be an enhanced stability of the newly formed place cells. By comparing the place cells formed in session 1 with the place cells formed in session 2 we could determine if modulating the activity of OLM interneurons lead to an increased retention of place cells encoding specific regions of the track. Neither stimulation nor inhibition of OLM interneurons resulted in *de novo* place cells that were more or less likely to be retained in the next session, nor did it lead to place cells that had altered spatial information content or place field widths (Figure 3J, K, Figure S2). However, of the place cells that did remain between sessions the degree of remapping to a new location on the track was altered. Enhancing the activity of OLM interneurons during place cell formation resulted in place cells that were more likely to encode the same location in subsequent sessions. When reducing the activity of OLM interneurons this resulted in a trend towards the formation of place cells that had a higher propensity to remap to a different location in subsequent sessions. (Figure 3J, Figure S3C,D).

Next, we analysed the stability of individual place cells between session 1 and session 2 in the novel environment. Increasing OLM activity during place cell formation led to a population of place cells with significantly higher spatial firing stability between the two sessions and place fields that shifted their location significantly less compared to place cells formed in the absence of increased OLM activity (0.363 ± 0.049 vs 0.634 ± 0.049 spatial correlation, 11.016 ± 2.854cm vs –4.231 ± 2.621cm place field shifts). Inhibiting OLM activity during place cell formation did not significantly alter the place cells spatial stability or alter the degree of place field location changes but there was a trend towards lower spatial firing stability and location shifting (Figure 3L, M). Therefore, OLM activity not only regulates the number of *de novo* place cells that are formed but also the population of place cells with high spatial stability. This suggests that increased OLM activity reduces the proportion of unstable place cells whilst reduced OLM activity leads to an increased proportion of unstable place cells. By categorising the place cells into stable and unstable based on a spatial correlation threshold of 0.5 we observe this to be the case with OLM activity bidirectionally altering the level of unstable place cells (Figure S3C,D).

Place cell representations completely remap in a novel environment but in the hippocampus there is also continual refinement of spatial representations even in familiar environments ^48^. To investigate whether OLM interneurons also regulate place cell stability in familiar environments or whether their role is selective for novel environments we next manipulated OLM activity during exploration of a familiar environment. In agreement with previous studies ^48^, the stability of place cell representations between sessions on a familiar track was notably higher than those of newly learnt representations but still resulted in measurable drift (Figure 4 A,C,F,G). Similar to the experiments in novel environments, OLM activity was selectively increased or decreased via optogenetic activation of ChrimsonR or JAWs on a defined section of the track. The acute effects of manipulating OLM activity were similar to that found in novel environments; increasing OLM activity reduced the number of place cells representing the stimulated region of the track whereas decreasing OLM activity had a limited effect with a trend towards an increased number of place cells (10% vs 21% and 16% vs 18%). This again demonstrates that although OLM interneurons have the capacity to inhibit place cells, at basal firing rates in familiar or novel conditions OLM interneurons do not appear to be actively inhibiting place cells.

**Figure 4.**
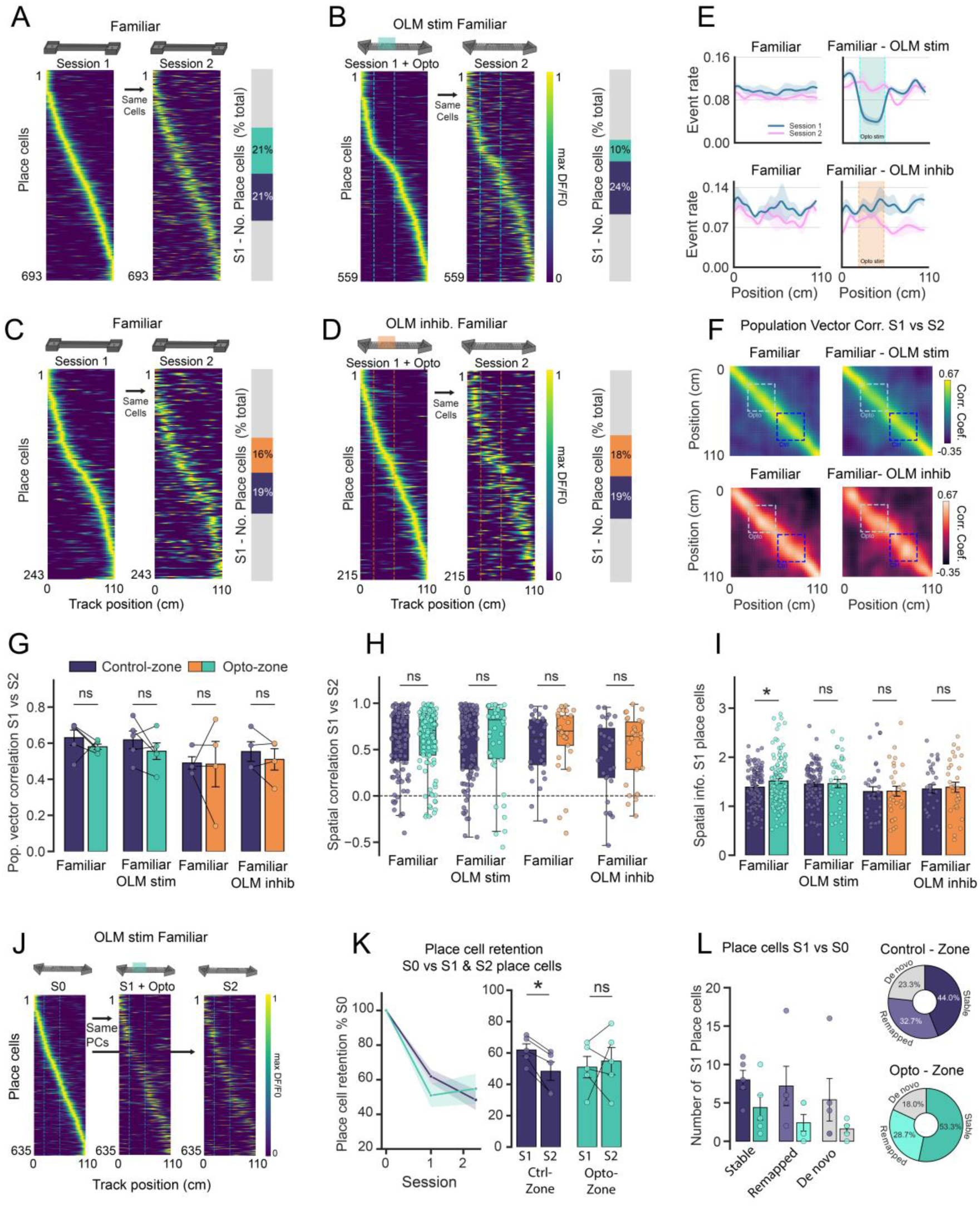
– Spatial representations in familiar environments are regulated by OLM activity. (A) Place cell firing rate maps in a familiar environment during session 1 in the absence of optogenetic stimulation of excitatory opsin ChrimsonR. Activity of the same cells in session 1 during a second exposure to the familiar environment (session 2) (left). Proportions of place cells during session 1 encoding different zones on the track. (B) Place cell firing rate maps in a familiar environment with optogenetic stimulation of excitatory opsin ChrimsonR at specific portion of the track (session 1). Activity of the same cells in session 1 during a second exposure to the familiar environment (session 2) (left). Proportions of place cells during session 1 encoding different zones on the track. (C) Same as A except for mice expressing the JAWs opsin in OLM interneurons. (D) Same as B except for optogenetic activation of JAWs to decrease OLM interneuron activity during familiar exploration. (E) Ca^2+^ event rate of all recorded neurons along the length of the track during the exploration of familiar environments (OLM stim (top row) n = 5 mice, OLM inhib. (bottom row) n = 4 mice) (F) Population vector correlation matrix between activity rate maps in session 1 vs session 2 for C-F, averaged across animals. (G) Average population vector correlation taken as average diagonal correlation for each track zone, (Control-zone vs Opto-zone). (OLM stim n = 5 mice, OLM inhib. n = 4 mice. (H) Spatial correlation values between place cells in session 1 and the activity of the same cells in session 2, for both control and optogenetic zones. (I) Spatial information content of session 1 place cells in the control and optogenetic zones. (paired t-test *: *p* < 0.05) (J) Place cell rate map retention between three exploitations of a familiar location (sessions 0-2) with OLM activity stimulation in middle session (session 1). Place cell rate maps from place cells in session 0 aligned to activity of the same cells that are also place cells in session 1 or session 2. (K) Quantification of J, number of place cells in session 1 and session 2 retained from session 0. (Paired t-test *: *p* < 0.05) (L) Place cell retention for place cells in session 1 vs same cells in session 0. Stable place cells: place cells in session 1 also a place cell in session 0 encoding the same location. Remapped place cells: place cells in session 1 also a place cell in session 0 but encoding a different location. *De novo* place cells: place cell in session 1 that was not a place cell in session 0. Percentage of, stable, remapped and *de novo* place cells in optogenetic stimulation and control zones.

In a novel environment, reducing OLM activity facilitates place cell remapping and increasing OLM activity promotes stability (Figure 3). In a familiar environment neither decreasing nor increasing OLM activity altered place cell stability, with no changes in the population vector or spatial correlations, alongside no changes in place cell information content or place field widths as a result of modulating OLM activity (Figure 4 F-I, Figure S2). These findings suggest that OLM interneuron regulation of place cell stability is contingent on the conditions found during novel explorations.

Although OLM activity did not alter place cell stability, enhancing their activity was able to reduce the number of place cells encoding the stimulated region of the track. In a familiar environment the reduction in place cell numbers may be due to silencing pre-existing place cells or reducing the rate of *de novo* place cell formation or remapping. To investigate the degree to which increased OLM activity may be silencing pre-existing place cells we measured the retention of pre-existing place cells from a previous exploration of the same familiar environment (session 0). We took the place cells in session 0 that had a place field centre in either the control or optogenetic stimulation zones and calculated the percentage of these place cells that were also a place cell in session 1 and session 2. For the control zone the number of session 0 place cells retained was sequentially reduced during session 1 and session 2, indicating after each exposure less session 0 place cells remained in the representation. However, during session 1, OLM stimulation led to a more drastic reduction in the number of session 0 place cells that were retained, to a comparable degree of place cell retention observed for session 2 (Figure 4 J,K; 61.91 ± 3.88% vs 48.34 ± 5.86% control zone, 50.96 ± 6.84% vs 54.81 ± 8.71% opto zone). This likely suggests that increased OLM activity during session 1 silenced a proportion of session 0 place cells that subsequently returned in session 2.

Increased OLM activity reduced the number of *de novo* place cells formed during exploration of a novel environment. To investigate if enhancing OLM activity also altered the degree of *de novo* place cell formation in a familiar environment we compared the place cells in session 1 to the same cells in session 0. We found that the percentage of *de novo* place cells (place cells in session 1 but not place cells in session 0), was only slightly reduced in the optogenetic stimulation zone compared to the control zone (18.03 ± 5.39% vs 23.31 ± 7.23%, opto vs control zone) suggesting that the role of OLM interneurons in modulating place cell formation is restricted to conditions in which the mouse is experiencing a novel environment (Figure 4L).

## Discussion

Synaptic plasticity and therefore remapping of place cells in CA1 is thought to be induced by large dendritic Ca^2+^ transients driven by strong input from temporoammonic fibers from entorhinal cortex coupled to input from CA3 ^6–15^. Here we show that OLM interneurons specifically inhibit this entorhinal input thereby reducing apical dendritic Ca^2+^ and synaptic plasticity during associative plasticity induced by pairing both inputs (Figure 1). This demonstrates the importance of OLM interneurons for gating behaviorally relevant synaptic plasticity. We then show that OLM activity is reduced at exactly the timepoint where synaptic plasticity is required to remap place cells – the experience of a novel environment (Figure 2). In a final set of experiments, we test the impact of manipulating OLM activity on place cell formation and stability in novel and familiar environments given the prediction that OLM inputs gate behaviorally relevant synaptic plasticity. These experiments show that artificially reinstating OLM activity in a novel environment reduces the emergence of new place cells but that the place cells that do emerge are more stable, in effect pruning out the more “speculative” place cells. In contrast, further reduction of OLM activity in a novel environment increased the proportion of unstable place cells (Figure 3 and S?). In a familiar environment, increasing OLM activity reduced place cell firing rates and the remaining place cells exhibited enhanced stability (Figure 4). For experiments examining place cell stability the use of control and OLM manipulation zones enabled us to make powerful within experiment comparisons. Overall, these results support the conclusion that OLM interneurons reduce synaptic plasticity and thereby stabilise place cells and spatial representations in the hippocampus. In a novel environment, OLM activity reduces to allow remapping of place cells.

We have previously demonstrated that inhibitory synapses from SOM interneurons (that include OLMα2 interneurons) to pyramidal neurons undergo potentiation when synaptic activation is paired with pyramidal neuron depolarization ^41^. By modelling the activity of place cells *in vivo* this inhibitory plasticity is predicted to regulate the stability of place cells in novel and familiar environments ^41, 47^. Interestingly, here we show that pairing of temporoammonic input with plateau potentials in CA1 pyramidal cells resulted in LTD of the TA input (Figure 1D) and therefore that plateau potentials cause a simultaneous increase in inhibition and decrease in excitation for synaptic transmission to distal CA1 dendrites. Under this framework the burst firing of place cells in their place fields drives potentiation of OLM synapses and depression of TA synapses which in turn makes further plasticity of those place cells less likely in response to ongoing place cell activity – effectively stabilizing the place cells. When OLM interneurons reduce their activity in novel environments less inhibitory plasticity is induced and therefore place cells can undergo more plasticity and exhibit less stability after remapping. This is indeed what we and other studies have found (c.f. Figure 3C and 4A) ^48–50^. Conversely, when we stimulate OLMα2 neurons in a novel environment there will be more inhibitory plasticity, at least in the place cells that are still active, making those place cells more stable. This is what we see in Figure 3 and therefore our results support these predictions for the role of inhibitory plasticity in place cell stabilisation.

An alternative explanation for the observed enhancement of place cell stability by OLM stimulation is that OLM activation specifically inhibits place cells that are more unstable leaving a population of stable place cells. Two pieces of data argue against this interpretation. Firstly, the proportion of stable place cells in the novel environment as a fraction of the total cells, and not just relative to unstable place cells, increases (Figure 3J). Secondly, in the familiar environment when place cells are compared between sessions prior to and during OLM stimulation, stable, unstable and novel place cells are all reduced equally in the stimulation and control zones of the track (Figure 4L).

The reduction in OLM activity when mice are placed in a novel environment is in line with reductions in SST interneuron activity in novel environments ^28–31^ suggesting either that OLM interneurons (and specifically OLMα2 neurons) form the majority of observed SST interneurons in CA1 or that a majority of SST interneurons respond in a similar manner to novelty. There is some evidence for both conclusions. The stratum oriens location of the OLM cell bodies makes them more accessible to imaging than deeper SST interneurons and therefore more likely to be recorded ^28, 46^ but where SST interneurons have been compared across different layers there does not appear to be a big difference in their response to novelty ^31^. A significant proportion of OLM interneurons do not express nicotinic a2 receptors and target their outputs more selectively to CA1 pyramidal neurons and not to fast spiking PV interneurons ^36^. There is also evidence that the proportion of non-OLMα2 to OLMα2 interneurons is higher in dorsal CA1 compared to ventral ^51^. We have not addressed the physiology of these non-OLMα2 OLM interneurons but the evidence suggests they respond to novelty and control dendritic excitability and synaptic plasticity in much the same way as OLMα2 interneurons. The increase in OLM activity with velocity of movement also mirrors findings from SST interneurons ^28, 29, 31, 46^ although there is some heterogeneity ^29, 46^. This heterogeneity might be explained by the heterogeneity of SST interneurons in general and OLM interneurons in particular ^36^ but we show that the reduced average velocity in novel environments does not account for the reduced OLM activity in response to novelty.

An important unanswered question is what controls the activity of OLM interneurons and specifically what causes reduced activity in novel environments? The main excitatory input to OLM interneurons is from CA1 pyramidal neurons but their average activity remains broadly stable in a novel environment so this is unlikely to cause the reduction of OLM activity. The major inhibitory inputs to OLM interneurons are from VIP expressing interneurons which strongly inhibit SST interneurons in both hippocampal and neocortical circuits ^17, 31, 52–56^. VIP expressing interneurons are themselves a diverse population of neurons that respond to inputs from hippocampal, cortical and subcortical areas and there are at least 2 subtypes that inhibit OLM interneurons. Interestingly, these subtypes are differentially modulated by animal velocity and therefore may be responsible for heterogeneity of OLM activity during movement^54^. Moreover, the activity of these VIP interneurons is increased in response to surprise or novelty, making them prime candidates to mediate the reduction of OLM activity with novelty ^31, 57, 58^. Then the question is what drives VIP interneurons and their activity increase in novel environments? 2 primary sources have been identified: Intercortical inputs and neuromodulatory inputs from subcortical structures. In the hippocampus these are for example from entorhinal cortex and cholinergic projections from medial septum, respectively ^17, 59–61^. This suggests dual antagonistic roles for cholinergic input to the circuit since direct input to OLM interneurons excites OLM interneurons whereas indirect input to VIP interneurons will inhibit OLM interneurons ^34, 38, 55, 62^. In a behavioral context it appears that the cholinergic input to VIP interneurons is dominant since cholinergic inputs are disinhibitory ^57, 63, 64^ and during novelty cholinergic inputs are active leading to a reduction in OLMα2 activity (Figure2) ^28, 30, 31^. However, during fear conditioning the aversive stimulus triggers cholinergic inputs and direct OLM activation ^34^.

SST interneurons and OLM interneurons specifically are critical for the learning of new contexts and when hippocampal OLM or SST interneurons are inhibited learning is impaired ^34, 37–39^. Overall, our data support a model where OLM activity is reduced in novel environments that require place cell representations to be remodelled by synaptic plasticity. The reduction in OLM activity is then a critical factor to provide a window for plasticity, and therefore place cell flexibility, that underpins learning.

## Acknowledgements

We thank Sarah Stuart for training in surgical procedures, Feng Xuan and Daniel Dombeck for advice with analysis and Klas Kullander for providing Chrna2-cre mice. We thank Claudia Clopath and David Dupret for input to conceptual development, Paul Chadderton, Peter Dayan and Daniel Dombeck for input to previous versions of the manuscript and all members of the Mellor group for discussion. This work was supported by Biotechnology and Biological Sciences Research Council (BBSRC, BB/N013956/1, BB/N019008/1), Wellcome Trust (101029/Z/13/Z, 108899/B/15/Z), Medical Research Council (MRC, MR/X010910/1).

## Contributions

Conceptualization, M.U. and J.R.M.; Methodology, M.U., M.C. and H-W. Z.; Investigation, M.U., M.C. and H-W. Z.; Visualization, M.U.; Writing – Original Draft, M.U. and J.R.M.; Writing – Review & Editing, M.U., M.C., H-W. Z. and J.R.M.; Funding Acquisition, M.U. and J.R.M.; Supervision, J.R.M.

## Declaration of interests

The authors declare no competing interests.

## Data availability

Data and code are available at https://github.com/mellor-lab/OLM_project_analysis and from the corresponding author.

## Methods

### Animal strains and breeding

All procedures and techniques were conducted in accordance with the UK animals scientific procedures act, 1986 with approval of the University of Bristol ethics committee. Chrna2-cre mice (Tg(Chrna2-cre)1Kldr) in which Cre recombinase is expressed under the nicotinic acetylcholine receptor α2 subunit (CHRNA2) promoter, was used to selectively target OLM interneurons ^33^. To express ChR2 or tdTomato within Chrna2 expressing interneurons C57/Bl6 homozygous Ai32 mice (Gt(ROSA)26Sor^tm32(CAG-COP4*H134R/EYFP)Hze^ Jax Stock number: 024109) or Ai14 mice (Gt(ROSA)26Sor^tm14(CAG-tdTomato)Hze^ Jax Stock number: 007908) were bred with Chrna2-cre mice creating heterozygous offspring with OLM specific expression of ChR2 or tdTomato. For *in vivo* experiments targeting OLM interneurons, Chrna2-cre mice were injected with Cre-specific viruses detailed below. Mice were group housed on a 12 hour light/dark cycle (lights on at 8am) for electrophysiological experiments and on a reversed 12 hour dark/light cycle (lights on at 8pm) for mice undergoing behavior experiments. Animal holding rooms were controlled with average ambient temperature of 21°C and 45% humidity. Male and female mice were used for all experiments.

### Slice preparation

Brain slices were prepared from mice aged 4 to 24 weeks following cervical dislocation and decapitation and brains removed and dissected in ice cold cutting solution (in mM: 205 Sucrose, 10 Glucose, 26 NaHCO_3_, 2.5 KCl, 1.25 NaH_2_PO_4_, 0.5 CaCl_2_, 5 MgSO_4_), continually bubbled with 95% O_2_ and 5% CO_2_. Sagittal brain slices, 400 µm thick containing the hippocampus were prepared via a VT1200 vibratome (Leica). Brain slices were transferred to aCSF (in mM: 124 NaCl, 3 KCl, 24 NaHCO_3_, 1.25 NaH_2_PO_4_ 10 Glucose, 2.5 CaCl_2_, 1.3 MgSO_4_), continually bubbled with 95% O_2_ and 5% CO_2._ and incubated at 35 °C for 30 min before being stored at room temperature for at least 30 min before use.

### Whole-cell patch clamp

Brain slices prepared from Chrna2-Cre x Ai32 animals were transferred to a submerged slice recording chamber with a constant 2.5 ml/min flow of aCSF, held at 32 °C. Slices were visualised using infrared DIC optics on an Olympus BX-50WI microscope for LTP experiments and a SliceScope Pro 6000/Multiphoton Imaging System (Scientifica) for 2photon imaging experiments. Patch electrodes with a resistance of 3-6 MΩ were pulled from borosilicate filamented glass capillaries (1.5 OD x 0.86 ID x 100 L mm, Harvard Apparatus) with a horizontal puller (P-97, Sutter Instrument Co., UK) and filled with internal solution.

Whole cell recordings were made with a MultiClamp 700A amplifier (Molecular Devices, USA), filtered at 2.4 kHz and digitised at 10 kHz with a CED Power1401 data acquisition board and Signal 5.12 acquisition software (CED, Cambridge, UK). Series resistance was monitored throughout all experiments and cells that showed >40% change were discarded from subsequent analysis. Recordings were also rejected from analysis if the series resistance was greater than 30 MΩ.

### Voltage clamp LTP recordings

Cells were voltage clamped at –70 mV to obtain excitatory currents. The internal pipette solution contained (in mM) 120 KMeSO_3_, 10 HEPES, 0.2 EGTA, 4 Mg-ATP, 0.3 Na-GTP, 8 NaCl, 10 KCl and adjusted to pH 7.4, 280-285 mOsm. No correction was made for the junction potential. Two Bipolar stimulating electrodes were placed in Stratum Radiatum (SR) and one in the Stratum Lacunosum Moleculare (SLM) to evoke independent excitatory synaptic responses from the Schaffer collateral (SC) and temporammonic (TA) pathways ^65^. The three independent pathways denoted: SC test pathway, TA test pathway and SC control pathway were stimulated separately at 0.05 Hz. A steady 5 min baseline of EPSC amplitudes was achieved before attempting to induce plasticity via the application of a theta burst stimulation (TBS) to the two test pathways. The TBS performed in current-clamp configuration consisted of a train of 10 bursts where each burst contained 5 pulses at 100 Hz with the frequency of bursts set at 5 Hz. The TBS was applied through the two test pathway stimulating electrodes either in combination with light pulses following the same pattern, to activate the ChR2-expressing OLM interneurons, or in the absence of light. Inhibitory synaptic responses during the TBS were evoked optogenetically via 2 ms square light pulse via a mounted 470 nm LED (Thorlabs, US) through a 40x objective lens.

LTP induction was performed within 10 min of whole-cell configuration to prevent plasticity washout. Following TBS, responses from all pathways were recorded for a further 25 min. Consecutive traces were averaged to produce a mean response every 30 seconds. The mean amplitude response of the baseline period was used to normalise the responses of each pathway. Plasticity was assessed by comparing the average ESPC amplitude during the last 5 min of the experiment between the control and test pathways.

### Two-photon Ca^2+^ imaging

For dendritic 2photon imaging experiments an internal solution containing (in mM:130 KMeSO_3_, 8 NaCl, 1 MgCl_2_, 10 HEPES, 4 MgATP, 0.3 Na_2_GTP, 5 QX-314) was supplemented with a Ca^2+^ fluorescent indicator (Fluo-5F, 200 μM) and a fluorescent dye (Alexa Fluor 594, 30 μM). Whole-cell recordings of CA1 pyramidal neurons established in voltage clamp (−70 mV) were then switched to current clamp and dye allowed to defuse into the neuron for at least 20 min. Bipolar stimulating electrodes were placed in the SR and SLM regions and used to evoke bursts of synaptic stimulation (5 pulses at 100 Hz). Secondary apical dendrites in both SR and SLM regions were imaged via a 60x objective lens (Olympus) with fluorescence excitation provided via a tuneable Ti:Sapphire pulsed laser (Newport Spectra-Physics) tuned to 810 nm. Dendrites were initially visualized in raster scanning mode and during synaptic stimulation individual dendritic branches were imaged via a line scanning mode each line scan consisting of 1,000 lines per second for 1 s. Synaptic stimulation and dendritic imaging was repeated 4 times for each dendrite with an interval of 20 sec with the resulting somatic voltage and dendritic fluorescent traces averaged across repeats. Images were acquired with a data acquisition board (National Instruments Corporation) using ScanImage software (version 3.8). For optogenetic activation of OLM interneurons during Ca^2+^ calcium imaging (5 pulses of 470 nm light coincident with electrical stimulation) was provided by a laser-diode (Doric lenses LDFLS_473/070) attached to 105 µm diameter fibre optic cannulae (Thorlabs CFMLC21L02) positioned over the SLM layer. During optical stimulation, optogenetic light was blocked from the PMT with a mechanical optical beam shutter (SHB1T, Thorlabs), positioned above the objective lens.

### Histology and Immunohistochemistry

Brains were fixed via cardiac perfusion of Phosphate buffered saline (PBS) followed by 4% Formaldehyde in PBS. Brains were removed and stored in PFA for 24hrs and then transferred to 30% sucrose PBS solution for 48 hours. 40 µm thick sagittal slices were then obtained via freezing microtome. Slices were first washed with PBS and then incubated in a blocking solution containing 5% donkey serum and 0.2% Triton X-100 for 60 min at room temperature. Slices were subsequently incubated at 4°C for 24 hours in blocking solution containing anti-SST (1:1000 Santa Cruz SC-7819) to stain for somatostatin and then incubated in blocking solution containing anti-goat Alexa Fluor 488 (1:500, Invitrogen Antibodies A-11055) for 2.5 hours at room temperature. Slices were then washed with PBS and mounted on microscope slides with 1:1000 DAPI staining for visualization, and images acquired using a widefield fluorescence microscope.

To visualise viral expression and lens implant locations Brains were fixed via cardiac perfusion with a high calcium Phosphate buffered saline solution (in mM: 137 NaCl, 2.7 KCl, 8.1 Na_2_HPO_4_, 1.47 KH_2_PO_4_, 0.49 MgCl*6H_2_0, 0.9 CaCl, pH 7.4) to activate GCaMP6 followed by 4% Formaldehyde in PBS. Brains were removed and stored in PFA for 24hrs before 100 um coronal brain slices were obtained via a VT1200 vibratome (Leica). Slices were washed in PBS containing 1:1000 DAPI mounted on microscope slides and imaged via a widefield fluorescence microscope.

### Viruses

For Ca^2+^ imaging of OLM interneurons Chrna2-cre mice were injected with a Cre dependent GCaMP6s (AAV1.Syn.Flex-GCaMP6s-WRPE-SV40, Addgene #100845 titre: 1.6 x 10^13^GC/ml). For Ca^2+^ imaging of pyramidal neurons and manipulation of OLM interneurons, Chrna2-cre mice were injected with GCaMP6f under the CAMKII promotor (AAV1-CAMKII-GCAMP6f-WRPE-SV40 Addgene #100834, titre: 1:10 – 2.8 x 10^13^ GC/ml), along with either Cre dependent redshifted excitatory opsins ChrimsonR (AAV5-Syn-Flex-rc[ChrimsonR-tdTomato] Addgene # 62723 titre: 8.5 x 1012 GC/ml), redshifted inhibitory opsin JAWs (AAV-CAG-FLEX-rc[Jaws-KGC-tdTomato-ER2], Addgene #84446 titre 7.9 x 10^12^ GC/ml constructed by the VVF, Zurich) or the control reporter mCherry (AAV5-hsyn-DIO-mcherry, Addgene #50459 titre: 1:4 – 7×10¹² GC/mL).

### *in vivo* surgical procedures

C57BL/6 Chrna2-cre mice aged 5-8 weeks, underwent 2 surgical procedures. First stereotaxic injections of adeno-associated viruses (AAV) were conducted under (1.5-4%) isoflurane and 0.05 mg/kg buprenorphine. Viruses were injected via a 35G NanoFil needle (WPI) into the right hippocampus CA1 (anteriorposterior: –1.9, mediolateral: +1.5 and dorsoventral: –1.5 relative to Bregma). Each injection had a volume of 600nl and injected at 50nl/min. The needle was left in place for 10 min post injection to allow viruses to diffuse from the injection site before removing the needle.

Mice were left a week to recover post viral injection surgery after which they underwent GRIN lens and baseplate implantation surgery conducted under (1.5-4%) isoflurane, 0.05 mg/kg buprenorphine and 5mg/kg Carprofen. First a 1.5 mm craniotomy was drilled above the viral injection site. Cortex above the hippocampus was aspirated under constant irrigation with ice cold aCSF (in mM: 150 NaCl, 2.5 KCl, 10 HEPES, 1 CaCl_2_, 1 MgCl_2_, pH 7.3). Upon reaching the mediolateral white matter striations a 1 mm diameter 4 mm length integrated lens and baseplate (ProView integrated lens, Inscopix) was slowly lowered on top of the hippocampus (anteroposterior: –2.0, mediolateral: +1.5 and dorsoventral: –1.15 relative to Bregma). The lens was fixed to the skull first with a layer of super-bond (Sun Medical) followed by surrounding bone cement containing gentamicin (CMW 1, DePuy synthesis). Mice were allowed to recover for at least 3 weeks post surgery before behavioural training.

### Linear track behaviour experiments

Water restricted mice (0.0375ml/g per day with a minimum 85% initial body weight) were trained to run back and forth along a 140cm linear track with a 6µl 10% sucrose reward delivered within a 15cm reward zone at the track ends. Three distinct linear tracks were used each with a unique texture and distinct local cues on the track walls. Distal cues were provided by two sets of distinct curtains surrounding each track, providing track polarity. Each experiment day consisted of three sessions. For familiar experiments, three consecutive sessions on a familiar track and for novel experiments, one session on a familiar track followed by two consecutive sessions on a novel track. Each 10 min session was separated by a 3 min home cage rest within an opaque box beside the track and familiar tracks were considered familiar after completion of at least five separate consecutive sessions on the same track. Post hoc tracking of the animal position was achieved by tracking the position of a miniscope mounted LED in custom MATLAB software. We then used the mouse position coordinates to calculate the mouse velocity which was then smoothed using a moving average filter with a width of 2 sec.

### *in vivo* Ca^2+^ imaging

Ca^2+^ imaging of hippocampal neurons was performed via 1-photon miniaturised microscopes (nVoke 2.0, Inscopix). The miniscope was attached to the baseplate and animals allowed to move freely along the linear track via custom counter balanced pully system. Ca^2+^ imaging was acquired via 455 nm excitation wavelength LED to activate the GCaMP calcium indicator. Recordings were obtained at resolution of 1280 x 800 pixels at a rate of 20 frames a second (20Hz), with a field of view of 1050 μm x 650 μm. For each animal the focal plane, LED intensity and gain imaging parameters were established prior to the first day of recording and kept constant throughout all experiments. Optogenetic stimulation was provided through the miniscope via a (620 nm) LED light. To activate the excitatory opsin ChrimsonR, A continual train of 5ms light pulses at 20hz were applied at a light intensity of 20 mW/mm^2^. For activation of inhibitory opsin JAWs a continual square pulse at light intensity 5 mW/mm^2^ was applied. Animal behaviour tracking was achieved via webcam (C920S, Logitech) positioned above the track controlled via custom written Python programme that enabled automated reward delivery and optogenetic triggering via a COM port connection to an Arduino Uno. The Arduino, relayed TTL pulses to the Inscopix DAQ board to trigger optogenetic stimulation to the miniscope and to two sperate solenoid valves to trigger reward delivery. Near real time tracking was achieved with a delay of 1 video frame (33 ms) which was accounted for via staggered optogenetic stimulation zone for each lap direction. Behavioural recordings were acquired at 30 fps and synchronised to the start and end of Ca^2+^ imaging sessions via a sync LED in the behaviour video field of view.

### Ca^2+^ recording imagine processing

Ca^2+^ recordings were analysed via the Inscopix data processing software (IDPS) and Python API (version 1.9.5). Raw video files were temporally and spatially down sampled to 10 Hz and 320×200 pixels and filtered with a Gaussian blur bandpass filter (high-pass cutoff: 0.5/pixel and low-pass cutoff 0.005/pixel). Each video frame was then motion corrected to an initial reference frame. Motion correction was restricted to a region of interest around the fluorescent field of view.

To obtain cell footprints of excitatory pyramidal neurons Constrained Nonnegative Matrix Factorisation for micro-endoscopic data (CNMFe) ^66^ was applied via the IDPS software. Hyperparameters for the CNMFe algorithm were selected for each animal to maximise cell separation and minimise over segmentation of cell footprints and were kept consistent for each field of view. Ca^2+^ transients were initially identified as Ca^2+^ signals that had a rising amplitude above 9 times the median absolute deviation of the trace (MAD) or a rate of decay equal to or slower than an exponential decay with a tau of 0.2. Cell footprints with a spatial component above 1, a signal to noise ratio less than 10 and an event rate of less than 0.005Hz were discarded from further analysis, any remaining footprints were then manually inspected to ensure they were accurately detecting neuronal signals. The resulting cell footprint traces were then deconvolved using the Online Active Set method to Infer Spikes (OASIS) algorithm applied via the IDPS software. We used a single order model and resulting spikes with an amplitude below a SNR of 6 were removed from the exported spike times used for further analysis. The OASIS algorithm approximates the number and timing of ‘spikes’ from the Ca^2+^ fluorescent traces, although this algorithm is not accurate in detecting ground truth action potential spike times ^67^, it acts as a more accurate approximation for temporal burst activity of pyramidal neurons compared to single Ca^2+^ transient time points.

For Ca^2+^ imaging of OLM interneurons cell footprints were manually identified and the resulting raw fluorescent traces extracted from the video. A baseline for each fluorescent trace was calculated by applying a 10-second sliding window and selecting the lowest 1st percentile of fluorescence values within each window as the baseline. This baseline was then used to calculate the ΔF/F_0_ for each interneuron trace.

### OLM activity

To obtain the lap-by-lap OLM interneuron activity, ΔF/F0 Ca^2+^ activity of OLM interneurons was thresholded to contain only activity when the mouse was running above 7cm/sec. For each interneuron an average activity per lap was obtained which was then averaged across neurons. Lap activities were taken from the last 10 laps of session 1 and first 8-10 laps of session 2. For average lap velocity this was taken as the average running velocity (velocity higher than 7cm/s) for each mouse.

### Place cell identification

Spike activity inferred from the recorded Ca^2+^ transients were assigned a track coordinate based on the mous position. Due to the directionality of place cell firing fields ^68^ activity during left and right traversals of the track were analysed separately and activity was only including for running epochs where the mous velocity exceeded 7cm/sec for a minimum of 10 cm. The two 15cm reward zones at the end of the track were also excluded from the analysis The resulting neuron spike activity was binned and averaged into 40 spatial bins giving a 2.75 cm bin width. This activity was divided by the mouse’s occupancy within the bin and gaussian filtered (σ = 1.5). The resulting rate maps were used to classify a neuron as a place cell by calculating the spatial information for each neuron.

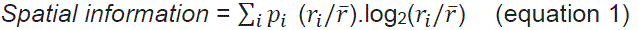

where *p*_*i*_ and r_*i*_ is probability and firing rate of the mouse/cell in position bin i and r̅ the mean firing rate.

This spatial information value was then compared to a distribution of spatial information values acquired by shuffling the spike times 1000 times. Statistically significant spatial coding was calculated using a z-test comparing the actual spatial information value to the bootstrapped spatial information distribution. For neurons that were classified as place cells for both left and right lap direction the rate map for the direction that had the lowest firing rate was discarded. All other rate maps for left and right traversals were normalised to the maximum firing bin and combined for all subsequent analysis. Place field centres were calculated as the bin with the maximal normalised activity. Place field bins are defined as the consecutive bins from the place field centre that has an activity rate equal to or exceeding 50% of event rate in the centre bin. Place field widths are defined as the sum of the bin widths in the place field. Cross-validation of place fields was performed on the initial familiar track dataset. Fields for even and odd laps were separated and sorted by even lap activity and showed high spatial and population vector correlation (Figure S3A,B).

### Population vector correlations

To compare the similarity of spatial representations across sessions, we calculated the population vector correlation between sessions. The population vector is defined as the activity rate for each neuron at a given positional bin. These population vectors are then correlated (using Pearson’s correlation) for each positional bin, resulting in a pairwise matrix of correlation values. For comparisons between sessions, the population vector activity is defined as the activity of place cells in session 1, with the same cells’ activity, irrespective of place cell status, in session 2. The population correlations for optogenetic and control zones are computed by taking the diagonal elements of the correlation matrix, corresponding to each zone, and calculated from the rate maps of each animal.

### Spatial correlation calculations

To compare the similarity between a cells spatial firing a spatial correlation for each cell is calculated as the Pearson’s correlation between the place cell activity in session 1 with the with the same cells’ activity, irrespective of place cell status, in session 2. If the neurons activity in session 2 is zero the spatial correlation is excluded from the analysis.

### Place cell retention

Place cell retention between sessions was calculated by taking the sorted place cells from the initial session and comparing them to same cells in subsequent sessions. Place cells were classed as lost if the place cell in the initial session was not classified as a place cell in subsequent sessions. Place cells were considered retained if the place cell in the initial session was also a place cell in subsequent sessions. These retained place cells were then split into stable place cells if the place field centres of the two cells were within 2 spatial bins and remapped place cells if the place field centres were more than 2 spatial bins apart. When comparing place cell retention to a previous session (Figure 4L) de novo place cells were classified as place cells in the subsequent session that were not place cells in the previous session.

### Statistical analysis

Experimental unit was defined by analysis of the source of greatest variance in the data. For *ex vivo* experiments and place cell and OLM interneuron *in vivo* measurements the experimental unit is cell and for behavioural experiments and population vector correlation analysis it is animal. Data in the text and in the figures are presented as mean ± SEM unless otherwise stated. The level of significance was assigned * if p < 0.05, ** if p < 0.01 and *** if p < 0.001 for statistical comparisons of all datasets. Data were processed and analysed using custom MATLAB and Python code.

## Supplementary Figures

**Figure S1.**
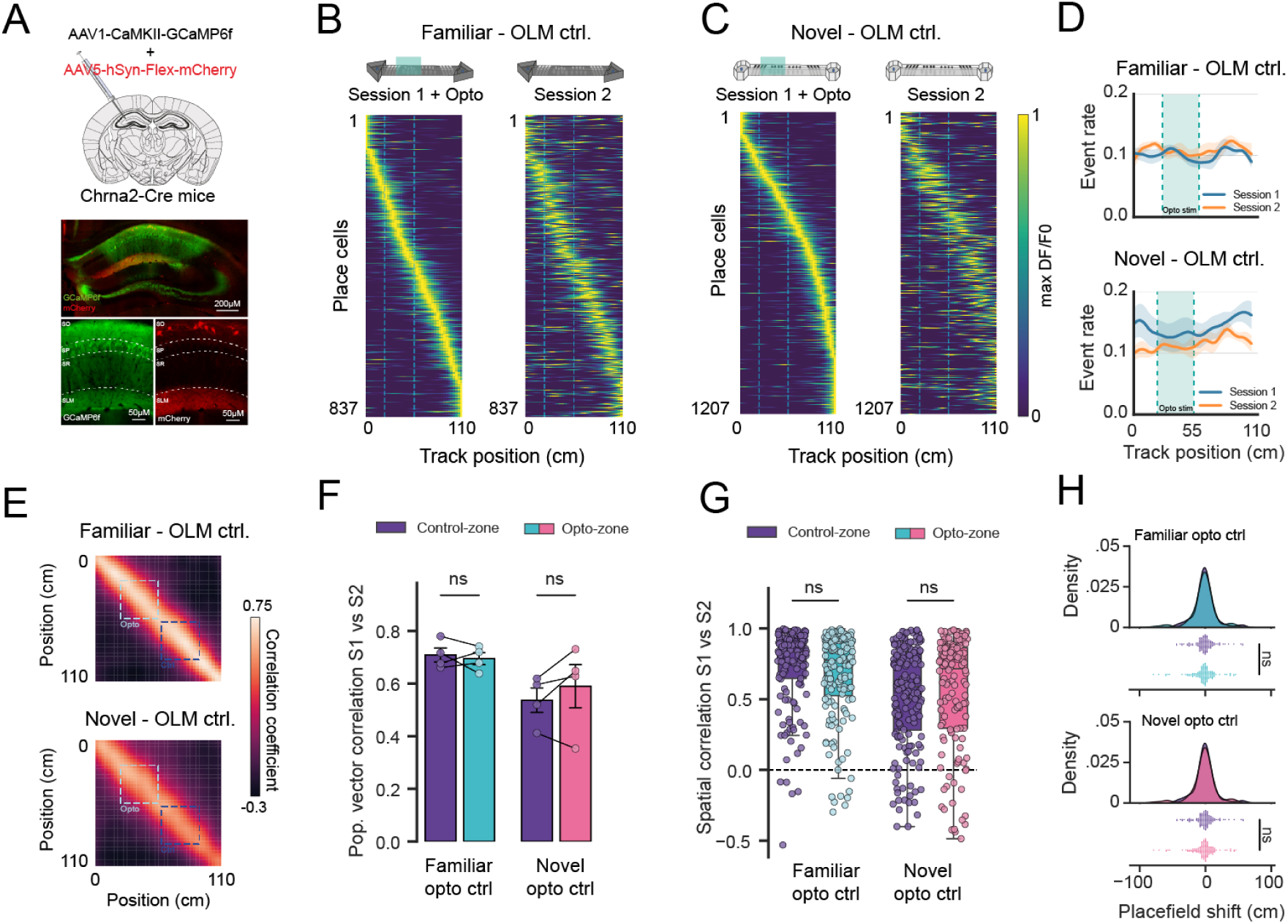
– mCherry control for optogenetic stimulation of OLM interneurons in familiar and novel environments. (A) Chrna2-cre mice were injected into the dorsal hippocampus with viral constructs to express GCaMP6f in pyramidal and mCherry into OLM interneurons. Example expression of mCherry in OLM interneurons. (B) Place cells acquired in a familiar environment (session 1) with light stimulation on opto zone. Activity of the same cells in session 1 during the second exposure to the familiar environment (session 2). (C) Same as B except for the first exposure to a novel environment. (D) Ca^2+^ event rate of all recorded neurons along the length of the track during the exploration of familiar or novel environments (Familiar n = 4 mice, Novel n = 4 mice) (E) Population vector correlation matrix between activity rate maps in session 1 vs session 2 for familiar and novel environments in B,C, averaged across animals. (F) Average population vector correlation taken as average diagonal correlation for each track zone, (Control-zone vs Opto-zone). (Familiar n = 4 mice, Novel n = 4 mice). (G) Spatial correlation values between place cells in session 1 and the activity of the same cells in session 2, for both control and optogenetic zones, in familiar and novel environments. (H) Place field location shifts from place cells in session 1 vs place field location of same cells in session 2.

**Figure S2.**
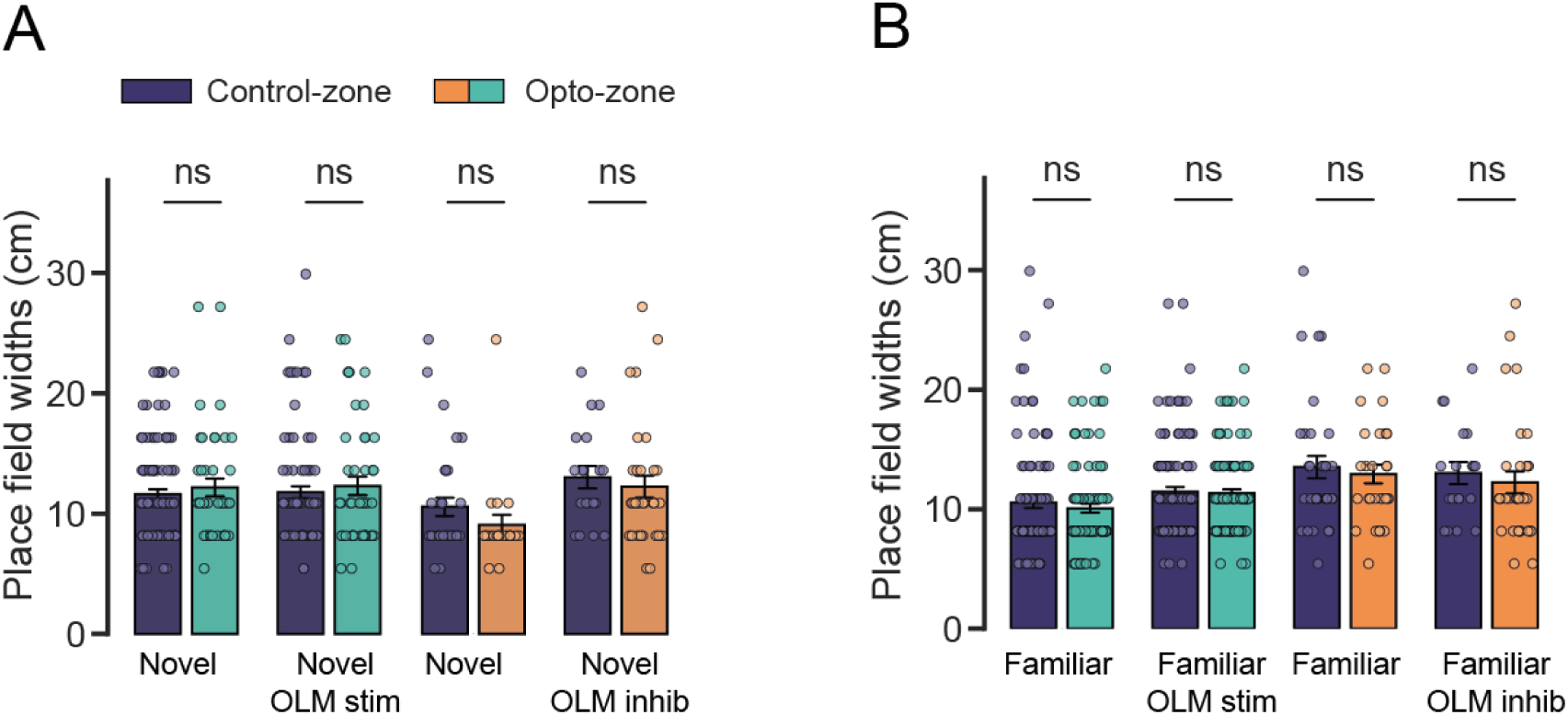
– No change in place field widths with OLM stimulation or inhibition in novel or familiar environments. Place field widths in control or opto stim zones for OLM stimulation or inhibition in novel (A) or familiar (B) environments.

**Figure S3.**
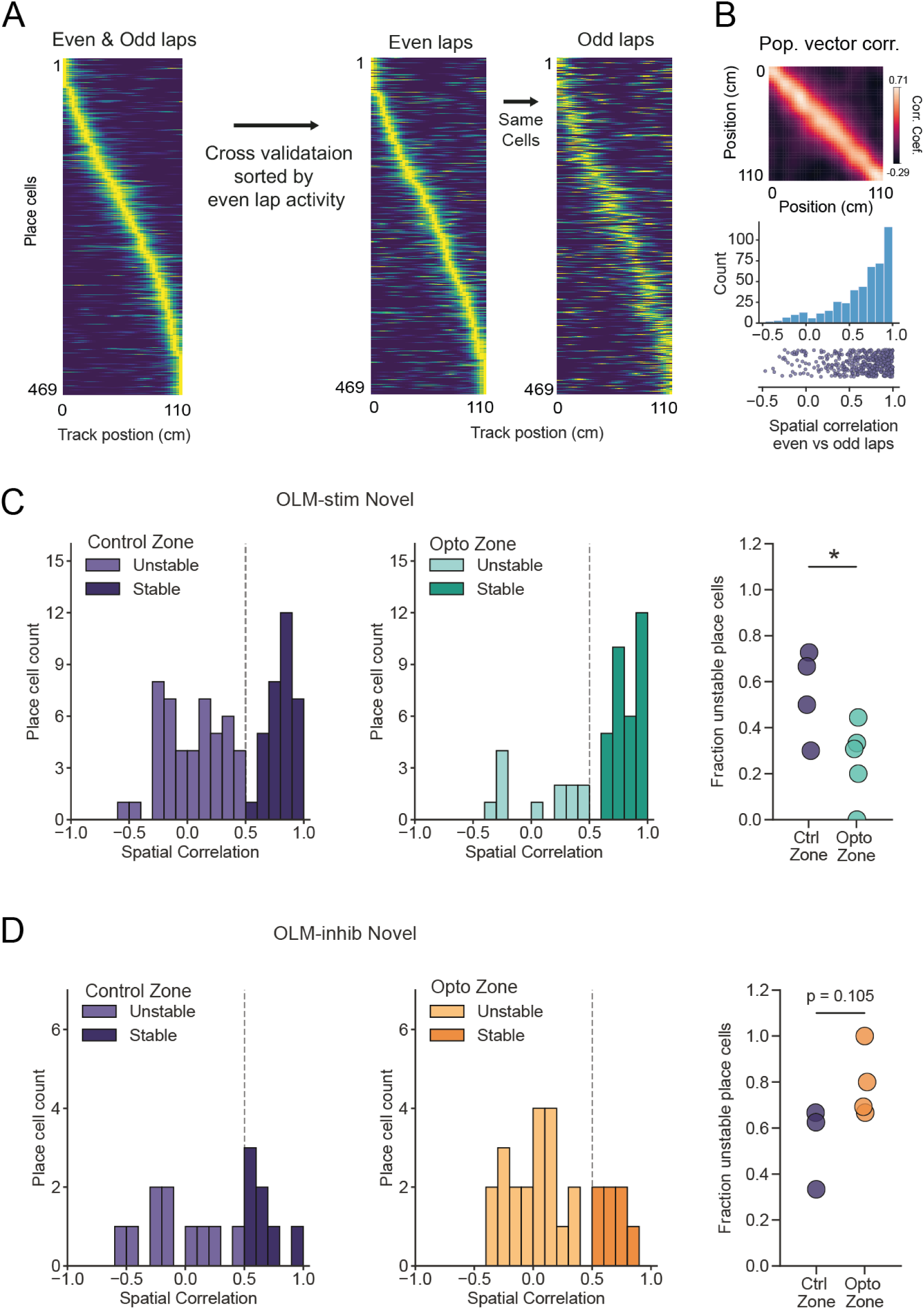
– Cross-validation of place fields and proportions of stable and unstable place cells after OLM stimulation or inhibition in a novel environment. (A) Average place cell population (left) split into even laps (middle) and odd maps sorted same as even laps (right) for mice in a familiar environment. (B) Population vector correlations (top) and spatial correlations (bottom) between even and odd laps. (C) Spatial correlation distribution for place cells in control and opto zones compared between sessions 1 and 2 for OLM stimulation in a novel environment. Classification of stable or unstable cells was set using a threshold of 0.5 spatial correlation. Proportion of unstable place cells for each mouse is plotted on the right. (D) Same as C but for OLM inhibition.

## Notes

### Competing Interest Statement

The authors have declared no competing interest.

## References

1. Fenton, A.A. Remapping revisited: how the hippocampus represents different spaces. Nat Rev Neurosci 25, 428–448 (2024).

2. Colgin, L.L., Moser, E.I. & Moser, M.B. Understanding memory through hippocampal remapping. Trends Neurosci 31, 469–477 (2008).

3. Dupret, D., O’Neill, J., Pleydell-Bouverie, B. & Csicsvari, J. The reorganization and reactivation of hippocampal maps predict spatial memory performance. Nat Neurosci 13, 995–1002 (2010).

4. Muller, R.U. & Kubie, J.L. The effects of changes in the environment on the spatial firing of hippocampal complex-spike cells. J Neurosci 7, 1951–1968 (1987).

5. Robinson, N.T.M., et al. Targeted Activation of Hippocampal Place Cells Drives Memory-Guided Spatial Behavior. Cell 183, 2041–2042 (2020).

6. Bittner, K.C., et al. Conjunctive input processing drives feature selectivity in hippocampal CA1 neurons. Nat Neurosci 18, 1133–1142 (2015).

7. Bittner, K.C., Milstein, A.D., Grienberger, C., Romani, S. & Magee, J.C. Behavioral time scale synaptic plasticity underlies CA1 place fields. Science 357, 1033–1036 (2017).

8. Magee, J.C. & Grienberger, C. Synaptic Plasticity Forms and Functions. Annu Rev Neurosci 43, 95–117 (2020).

9. Grienberger, C. & Magee, J.C. Entorhinal cortex directs learning-related changes in CA1 representations. Nature 611, 554–562 (2022).

10. Priestley, J.B., Bowler, J.C., Rolotti, S.V., Fusi, S. & Losonczy, A. Signatures of rapid plasticity in hippocampal CA1 representations during novel experiences. Neuron 110, 1978–1992 e1976 (2022).

11. Diamantaki, M., et al. Manipulating Hippocampal Place Cell Activity by Single-Cell Stimulation in Freely Moving Mice. Cell reports 23, 32–38 (2018).

12. Sheffield, M.E. & Dombeck, D.A. Calcium transient prevalence across the dendritic arbour predicts place field properties. Nature 517, 200–204 (2015).

13. Stuart, S.A., et al. Hippocampal-dependent navigation in head-fixed mice using a floating real-world environment. Scientific reports 14, 14315 (2024).

14. Takahashi, H. & Magee, J.C. Pathway interactions and synaptic plasticity in the dendritic tuft regions of CA1 pyramidal neurons. Neuron 62, 102–111 (2009).

15. Golding, N.L., Staff, N.P. & Spruston, N. Dendritic spikes as a mechanism for cooperative long-term potentiation. Nature 418, 326–331 (2002).

16. Booker, S.A. & Vida, I. Morphological diversity and connectivity of hippocampal interneurons. Cell Tissue Res 373, 619–641 (2018).

17. Topolnik, L. & Tamboli, S. The role of inhibitory circuits in hippocampal memory processing. Nat Rev Neurosci 23, 476–492 (2022).

18. Klausberger, T. & Somogyi, P. Neuronal diversity and temporal dynamics: the unity of hippocampal circuit operations. Science 321, 53–57 (2008).

19. Pelkey, K.A., et al. Hippocampal GABAergic Inhibitory Interneurons. Physiol Rev 97, 1619–1747 (2017).

20. Freund, T.F. & Buzsaki, G. Interneurons of the hippocampus. Hippocampus 6, 347–470 (1996).

21. Harris, K.D., et al. Classes and continua of hippocampal CA1 inhibitory neurons revealed by single-cell transcriptomics. PLoS biology 16, e2006387 (2018).

22. Rolotti, S.V., et al. Local feedback inhibition tightly controls rapid formation of hippocampal place fields. Neuron 110, 783–794 e786 (2022).

23. Grienberger, C., Milstein, A.D., Bittner, K.C., Romani, S. & Magee, J.C. Inhibitory suppression of heterogeneously tuned excitation enhances spatial coding in CA1 place cells. Nat Neurosci 20, 417–426 (2017).

24. Valero, M., Navas-Olive, A., de la Prida, L.M. & Buzsaki, G. Inhibitory conductance controls place field dynamics in the hippocampus. Cell reports 40, 111232 (2022).

25. Royer, S., et al. Control of timing, rate and bursts of hippocampal place cells by dendritic and somatic inhibition. Nat Neurosci 15, 769–775 (2012).

26. Milstein, A.D., et al. Inhibitory Gating of Input Comparison in the CA1 Microcircuit. Neuron 87, 1274–1289 (2015).

27. Katona, L., et al. Sleep and movement differentiates actions of two types of somatostatin-expressing GABAergic interneuron in rat hippocampus. Neuron 82, 872–886 (2014).

28. Sheffield, M.E.J., Adoff, M.D. & Dombeck, D.A. Increased Prevalence of Calcium Transients across the Dendritic Arbor during Place Field Formation. Neuron 96, 490–504 e495 (2017).

29. Hainmueller, T., Cazala, A., Huang, L.W. & Bartos, M. Subfield-specific interneuron circuits govern the hippocampal response to novelty in male mice. Nature communications 15, 714 (2024).

30. Arriaga, M. & Han, E.B. Structured inhibitory activity dynamics in new virtual environments. Elife 8 (2019).

31. Geiller, T., et al. Large-Scale 3D Two-Photon Imaging of Molecularly Identified CA1 Interneuron Dynamics in Behaving Mice. Neuron 108, 968–983 e969 (2020).

32. Gu, Z. & Yakel, J.L. Timing-dependent septal cholinergic induction of dynamic hippocampal synaptic plasticity. Neuron 71, 155–165 (2011).

33. Leao, R.N., et al. OLM interneurons differentially modulate CA3 and entorhinal inputs to hippocampal CA1 neurons. Nat Neurosci 15, 1524–1530 (2012).

34. Lovett-Barron, M., et al. Dendritic inhibition in the hippocampus supports fear learning. Science 343, 857–863 (2014).

35. Hilscher, M.M., Mikulovic, S., Perry, S., Lundberg, S. & Kullander, K. The alpha2 nicotinic acetylcholine receptor, a subunit with unique and selective expression in inhibitory interneurons associated with principal cells. Pharmacol Res 196, 106895 (2023).

36. Chamberland, S., et al. Functional specialization of hippocampal somatostatin-expressing interneurons. Proc Natl Acad Sci U S A 121, e2306382121 (2024).

37. Siwani, S., et al. OLMalpha2 Cells Bidirectionally Modulate Learning. Neuron 99, 404–412 e403 (2018).

38. Haam, J., Zhou, J., Cui, G. & Yakel, J.L. Septal cholinergic neurons gate hippocampal output to entorhinal cortex via oriens lacunosum moleculare interneurons. Proc Natl Acad Sci U S A 115, E1886–E1895 (2018).

39. Szonyi, A., et al. Brainstem nucleus incertus controls contextual memory formation. Science 364 (2019).

40. Mikulovic, S., Restrepo, C.E., Hilscher, M.M., Kullander, K. & Leao, R.N. Novel markers for OLM interneurons in the hippocampus. Front Cell Neurosci 9, 201 (2015).

41. Udakis, M., Pedrosa, V., Chamberlain, S.E.L., Clopath, C. & Mellor, J.R. Interneuron-specific plasticity at parvalbumin and somatostatin inhibitory synapses onto CA1 pyramidal neurons shapes hippocampal output. Nature communications 11, 4395 (2020).

42. Griesius, S., et al. Reduced expression of the psychiatric risk gene DLG2 (PSD93) impairs hippocampal synaptic integration and plasticity. Neuropsychopharmacology (2022).

43. O’Dell, T.J. Behavioral Timescale Cooperativity and Competitive Synaptic Interactions Regulate the Induction of Complex Spike Burst-Dependent Long-Term Potentiation. J Neurosci 42, 2647–2661 (2022).

44. Stuart, G.J. & Spruston, N. Dendritic integration: 60 years of progress. Nat Neurosci 18, 1713–1721 (2015).

45. Williams, S.R. & Mitchell, S.J. Direct measurement of somatic voltage clamp errors in central neurons. Nat Neurosci 11, 790–798 (2008).

46. Arriaga, M. & Han, E.B. Dedicated Hippocampal Inhibitory Networks for Locomotion and Immobility. J Neurosci 37, 9222–9238 (2017).

47. Pedrosa, V. & Clopath, C. The interplay between somatic and dendritic inhibition promotes the emergence and stabilization of place fields. PLoS computational biology 16, e1007955 (2020).

48. Geva, N., Deitch, D., Rubin, A. & Ziv, Y. Time and experience differentially affect distinct aspects of hippocampal representational drift. Neuron 111, 2357–2366 e2355 (2023).

49. Ziv, Y., et al. Long-term dynamics of CA1 hippocampal place codes. Nat Neurosci 16, 264–266 (2013).

50. Hainmueller, T. & Bartos, M. Parallel emergence of stable and dynamic memory engrams in the hippocampus. Nature 558, 292–296 (2018).

51. Mikulovic, S., et al. Ventral hippocampal OLM cells control type 2 theta oscillations and response to predator odor. Nature communications 9, 3638 (2018).

52. Ren, C., et al. Global and subtype-specific modulation of cortical inhibitory neurons regulated by acetylcholine during motor learning. Neuron 110, 2334–2350 e2338 (2022).

53. Tyan, L., et al. Dendritic inhibition provided by interneuron-specific cells controls the firing rate and timing of the hippocampal feedback inhibitory circuitry. J Neurosci 34, 4534–4547 (2014).

54. Francavilla, R., et al. Connectivity and network state-dependent recruitment of long-range VIP-GABAergic neurons in the mouse hippocampus. Nature communications 9, 5043 (2018).

55. Williams, L.E. & Holtmaat, A. Higher-Order Thalamocortical Inputs Gate Synaptic Long-Term Potentiation via Disinhibition. Neuron 101, 91–102 e104 (2019).

56. Pfeffer, C.K., Xue, M., He, M., Huang, Z.J. & Scanziani, M. Inhibition of inhibition in visual cortex: the logic of connections between molecularly distinct interneurons. Nat Neurosci 16, 1068–1076 (2013).

57. Fu, Y., et al. A cortical circuit for gain control by behavioral state. Cell 156, 1139–1152 (2014).

58. Szadai, Z., et al. Cortex-wide response mode of VIP-expressing inhibitory neurons by reward and punishment. Elife 11 (2022).

59. Bell, L.A., Bell, K.A. & McQuiston, A.R. Activation of muscarinic receptors by ACh release in hippocampal CA1 depolarizes VIP but has varying effects on parvalbumin-expressing basket cells. J Physiol 593, 197–215 (2015).

60. Bell, L.A., Bell, K.A. & McQuiston, A.R. Acetylcholine release in mouse hippocampal CA1 preferentially activates inhibitory-selective interneurons via alpha4beta2* nicotinic receptor activation. Front Cell Neurosci 9, 115 (2015).

61. Luo, X., et al. Transcriptomic profile of the subiculum-projecting VIP GABAergic neurons in the mouse CA1 hippocampus. Brain Struct Funct 224, 2269–2280 (2019).

62. Bell, L.A., Bell, K.A. & McQuiston, A.R. Synaptic muscarinic response types in hippocampal CA1 interneurons depend on different levels of presynaptic activity and different muscarinic receptor subtypes. Neuropharmacology 73, 160–173 (2013).

63. Letzkus, J.J., et al. A disinhibitory microcircuit for associative fear learning in the auditory cortex. Nature 480, 331–335 (2011).

64. Pi, H.J., et al. Cortical interneurons that specialize in disinhibitory control. Nature 503, 521–524 (2013).

65. Palacios-Filardo, J., et al. Acetylcholine prioritises direct synaptic inputs from entorhinal cortex to CA1 by differential modulation of feedforward inhibitory circuits. Nature communications 12, 5475 (2021).

66. Zhou, P., et al. Efficient and accurate extraction of in vivo calcium signals from microendoscopic video data. Elife 7 (2018).

67. Pachitariu, M., Stringer, C. & Harris, K.D. Robustness of Spike Deconvolution for Neuronal Calcium Imaging. J Neurosci 38, 7976–7985 (2018).

68. Battaglia, F.P., Sutherland, G.R. & McNaughton, B.L. Local sensory cues and place cell directionality: additional evidence of prospective coding in the hippocampus. J Neurosci 24, 4541–4550 (2004).

